# The Trend of Suicide Reporting in the Media: an Effectiveness Study of Daily Surveillance over Nine Years

**DOI:** 10.1101/768945

**Authors:** Chia-Yi Wu, Ming-Been Lee, Shih-Cheng Liao, Chia-Ta Chan, Chun-Ying Chen

## Abstract

Inadequate suicide reports can cause copycat suicide attempts, but legal regulations for the media are rarely seen worldwide. In Taiwan, daily media surveillance with immediate feedback to correct inappropriate reports has been an interactive policy in the national suicide prevention strategy for a decade. We aimed to evaluate the effectiveness of current surveillance program via assessing the adherence rates with 12-item WHO responsible reporting guideline in print newspapers (2010-2018) and online media (2017-2018). The results showed that media reporting of suicide significantly improved in most guideline items under surveillance. But the development of psychiatric-media liaisons should further improve reporting quality.

## Introduction

Suicide is a major public health issue worldwide. To reduce suicide risk and enhance protective factors, well-established suicide prevention strategies comprise three levels: 1) universal strategy (education about suicide, means control, good quality media reporting and mental health promotion for the general public); 2) selective strategy (early identification, prevention and intervention for high-risk individuals); and 3) indicated strategy (surveillance and aftercare for suicide attempters) [1]. As one of the universal strategies, media reporting on suicide is a focus of major attention; it could serve as a risk or protective factor for suicide behavior. The media tend to report suicide in an exaggerated and overly detailed manner [2]. A number of studies have shown that inappropriate reporting has a negative impact on suicide [3,4]. The repetition of suicide reports, reporting on the front page, highlighting the details of suicide and glorifying suicide behaviors could increase the number of subsequent suicide acts in vulnerable populations [5]. In particular, inappropriate media reporting of celebrities or particular cases could lead to an accompanying increase in suicide rates [2-5, 6-9]. For example, recent detailed media reports and subsequent Google searches on carbon-monoxide self-poisoning (or charcoal-burning suicide) played a significant role in the huge increase in suicidal death by this method in East Asia [10-12]. To improve the quality of suicide reporting, the WHO released guidelines with a list of 6 “Dos” and 6 “Don’ts” for media professionals [13]. Some researchers have reported that implementation of WHO reporting guidelines could decrease the negative impact of suicide [14,15]. Fu et al. (2008) investigated 5,740 reports of suicide and found no significant changes in the content of reporting after publication of the WHO recommendations [16]. Chiang et al. analyzed the trends in major newspapers violating reporting recommendations; they found that suicide was the most commonly reported headline on the front page and on the whole page, with many reports including photographs [17]. Few studies have comprehensively analyzed compliance with the WHO guidelines, or examined the effectiveness of a long-term intervention on the media report of suicide. The nine-year observational study aimed to assess the effectiveness of a surveillance with interactive intervention on the quality of media suicide reporting, conducted by the Taiwan Suicide Prevention Center (TSPC).

## Method

### Study design

The study was ethically approved by Institutional Review Board where the corresponding author affiliated (M04B3020). This was a long-term observational study nested in a nine-year media observation program targeting at developing the national suicide prevention strategy in Taiwan. Media reporting of suicide has been under surveillance by trained staffs of the TSPC and the researchers for nearly a decade. In 2006, the TSPC published and promoted the Chinese version of the WHO guidelines for responsible media reporting of suicides; since 2010, the approach of regular surveillance included daily monitoring of media reporting on suicide and giving timely feedback to the media in order to improve quality of reports. Moreover, to enhance quality suicide reporting, the TSPC began an additional interactive approach in 2014 which includes: 1) a site visit with problem-based discussion for each newspaper company at the beginning of the year and 2) an annual liaison meeting involving media professionals, mental health specialists, legislators and government representatives, to enhance the reporting quality based on the data of suicide reporting in the previous six months. The researchers evaluated the longitudinal effects of two media surveillance approaches (i.e., regular surveillance vs. interactive surveillance) from two major report platforms (i.e. paper or online).

### Data Sources and the coding

Four major newspaper companies with the highest readership rates in Taiwan were selected for the study: China Times, Liberty Times, Apple Daily and United Daily News. All four newspapers offer both print and online platforms of reports. The data were prospectively collected from print newspapers and online news from 2010 to 2018. For print newspapers, we examined the content of all pages related to suicide (including completed suicides, suicide attempts and suspected suicides) and marked them for coding. For online sources, we analyzed data for the years 2017 and 2018.

The coding for quantity analysis for print newspaper was based on the number of articles reporting suicide events. For instance, an incident event could be reported in different articles; the data were counted for all independent articles with a unique title. However, for online news, the coding was based on the number of suicide person-events covered. For example, an incident was counted as just one item even if more than one article reported the same incident. The sources of data might be limited to available news reports from difference platforms, but bias could be lowered since the TSPC has made efforts in collecting information and checked the reliability of the data. In addition, every piece of coding was rechecked through weekly meetings of the staffs and the researchers with cautions to avoid biased analysis and missing values.

### Data Management

Management of newspaper and online news text was conducted by a team of well-trained staff at the TSPC who were familiar with the definitions of the 12 WHO Dos and Don’ts for suicide reporting [13]. Coding of the suicide reports was done according to a standardized coding manual developed by the TSPC. The “What to Do” items (6 Dos) included: 1) work closely with health authorities in presenting the facts; 2) refer to suicide as completed, not successful; 3) present relevant data only on the inside pages; 4) highlight alternatives to suicide; 5) provide information on helplines and community resources; and 6) publicize risk indicators and warning signs. The “What Not to Do” items (6 Don’ts) included: 1) Do not publish photographs or suicide notes; 2) Do not report specific details of the method used; 3) Do not give simplistic reasons; 4) Do not glorify or sensationalize suicide; 5) Do not use religious or cultural stereotypes; and 6) Do not apportion blame.

We filtered and printed the news items meeting the inclusion criteria for coding. Reliability analysis of the assessors was then carried out based on the preliminary coding. When the coding items were difficult to define, all assessors discussed the item, under supervision by a senior leader at TSPC, until consensus was reached. All the recruited articles were coded for the 12 WHO recommendation items (Do/Don’t) as “yes” (adherence to the recommendation, coding 1) or “no” (violation of the recommendation, coding 0).

### Data Analysis

Data were analyzed through available sources of news reports in paper or online forms from four major newspaper companies. Descriptive statistics was expressed as the frequency of adherence to each WHO guideline. The Chi-squared test was used to analyze the trends of adherence to six Don’ts and six Dos in reporting suicide for four print newspapers from 2010 to 2018, for online news during 2017-2018 and to test the effectiveness of the interactive approach after 2014, using SPSS version 21 software (IBM Corp., Armonk, NY). In addition, adherence was also compared between the online and print media in 2017-2018.

## Results

### Adherence to the WHO suicide reporting guidelines for print newspapers

In total, 5,529 articles from print media and 16,445 person-event items from online media for the four major newspapers were selected for quantity analysis. Overall, the yearly number of suicide news reports across the nine years markedly decreased by 76.2% for the four major print newspapers, but the trend for online media sharply increased by 100% over the years (Supplementary Figure 1). As Table 1 shows, the annual adherence rates of print newspapers to the WHO guidelines differed significantly different for all items across the nine years. Among the Don’t guidelines (Table 1, Supplementary Figure 2), the following three items had steadily higher adherence rates (around 95%) over the years: “Do not glorify or sensationalize,” “Do not apportion blame” and “No religious or cultural stereotypes.” However, lower adherence was noted for the items “No photographs” (26.7-41.7%; average 35.0%) and “No simplistic suicide reasoning” (21.6-54.1%; average 34.1%). Importantly, adherence to the recommendations against reporting specific details of the suicide method and reporting simplistic suicide reasoning increased steadily from 2012 to 2018.

**Table 1.** The annual rates of adherence to WHO recommended guide1ines for four major print newspapers from 2010 to 2018.

|  | 2010 | 2011 | 2012 | 2013 | 2014 | 2015 | 2016 | 2017 | 2018 | Total |  |
| --- | --- | --- | --- | --- | --- | --- | --- | --- | --- | --- | --- |
| n (%) | (n=1136) | (n=829) | (n=881) | (n=672) | (n=563) | (n=499) | (n=381) | (n=298) | (n=270) | (N=5529) | X<sup>2</sup> |
| N1. Do not glorify or sensationalize | 1114 (98.1) | 803 (96.9) | 854 (96.9) | 664 (98.8) | 558 (99.1) | 473 (94.8) | 373 (97.9) | 295 (99.0) | 269 (99.6) | 5403 (97.7) | 40.1*** |
| N2. No religious or cultural stereotypes | 1108 (97.5) | 811 (97.8) | 840 (95.4) | 664 (98.8) | 557 (98.9) | 476 (95.4) | 364 (95.5) | 292 (98.0) | 263 (97.4) | 5375 (97.2) | 36.2*** |
| N3. Do not apportion blame | 970(85.4) | 760 (91.7) | 801 (90.9) | 629 (93.6) | 519 (92.2) | 426 (85.4) | 360 (94.5) | 263 (88.3) | 263 (97.4) | 4991 (90.3) | 82.4*** |
| N4. No specific details | 634 (55.8) | 468 (56.5) | 411 (46.7) | 384 (57.1) | 381 (67.7) | 339 (67.9) | 239 (62.7) | 220 (73.8) | 241 (89.3) | 3317 (60.0) | 228.5*** |
| N5. No photos | 432 (38.0) | 346 (41.7) | 357 (40.5) | 217 (32.3) | 173 (30.7) | 142 (28.5) | 117 (30.7) | 81 (27.2) | 72 (26.7) | 1937 (35.0) | 68.3*** |
| N6. No simplistic reasons | 413 (36.4) | 317 (38.2) | 190 (21.6) | 180 (26.8) | 159 (28.2) | 157 (31.5) | 175 (45.9) | 149 (50.0) | 146 (54.1) | 1886 (34.1) | 201.8*** |
| D1. Refer to completed not successful suicides | 1086 (95.6) | 751 (90.6) | 788 (89.4) | 642 (95.5) | 557 (98.9) | 486 (97.4) | 353 (92.7) | 291 (97.7) | 268 (99.3) | 5222 (94.4) | 119.9*** |
| D2. Present relevant data only on inside pages | 1063 (93.6) | 773 (93.2) | 788 (89.4) | 617 (91.8) | 497 (88.3) | 372 (74.6) | 332 (87.1) | 266 (89.3) | 259 (95.9) | 4967 (89.8) | 174.3*** |
| D3. Provide helplines information | 213 (18.8) | 216 (26.1) | 370 (42.0) | 381 (56.7) | 374 (66.4) | 353 (70.7) | 292 (76.6) | 233 (78.2) | 247 (91.5) | 2679 (48.5) | 1199.7*** |
| D4. Publicize risk indicators | 264 (23.2) | 292 (35.2) | 194 (22.0) | 193 (28.7) | 157 (27.9) | 188 (37.7) | 159 (41.7) | 57 (19.1) | 28 (10.4) | 1532 (27.7) | 162.9*** |
| D5. Discuss with experts | 142 (12.5) | 76 (9.2) | 119 (13.5) | 86 (12.8) | 78 (13.9) | 135 (27.1) | 70 (18.4) | 57 (19.1) | 34 (12.6) | 797 (14.4) | 99.6*** |
| D6. Highlight alternatives | 39 (3.4) | 38 (4.6) | 51 (5.8) | 35 (5.2) | 26 (4.6) | 49 (9.8) | 27 (7.1) | 31 (10.4) | 19 (7.0) | 315 (5.7) | 44.6*** |

Note: \*p<0.05 \*\*p<0.01 \*\*\*p<0.001

Regarding the six Dos (Table 1, Supplementary Figure 3), “Refer to completed, not successful suicides” (89.4-99.3%, average 94.4%) and “Publish relevant data only on inside pages” (74.6-95.9%, with average 89.8%) sustained the highest level of adherence over the years. There was a trend of marked increase in adherence to the item “Provide helpline information,” from 18.8% in 2010 to 91.1% in 2018. Most importantly, the following three items had persistently low adherence over the years: “Highlight alternatives” (average 5.7%), “Discuss with experts” (average 14.4%) and “Publicize risk indicators” (average 27.7%). Comparison of the adherence between phase 1 (prior to interactive approach) and phase 2 (after interactive approach) indicated that a majority of the items except for “No photograph” had significantly better adherence in phase 2 than in phase 1 (Table 2).

**Table 2:**
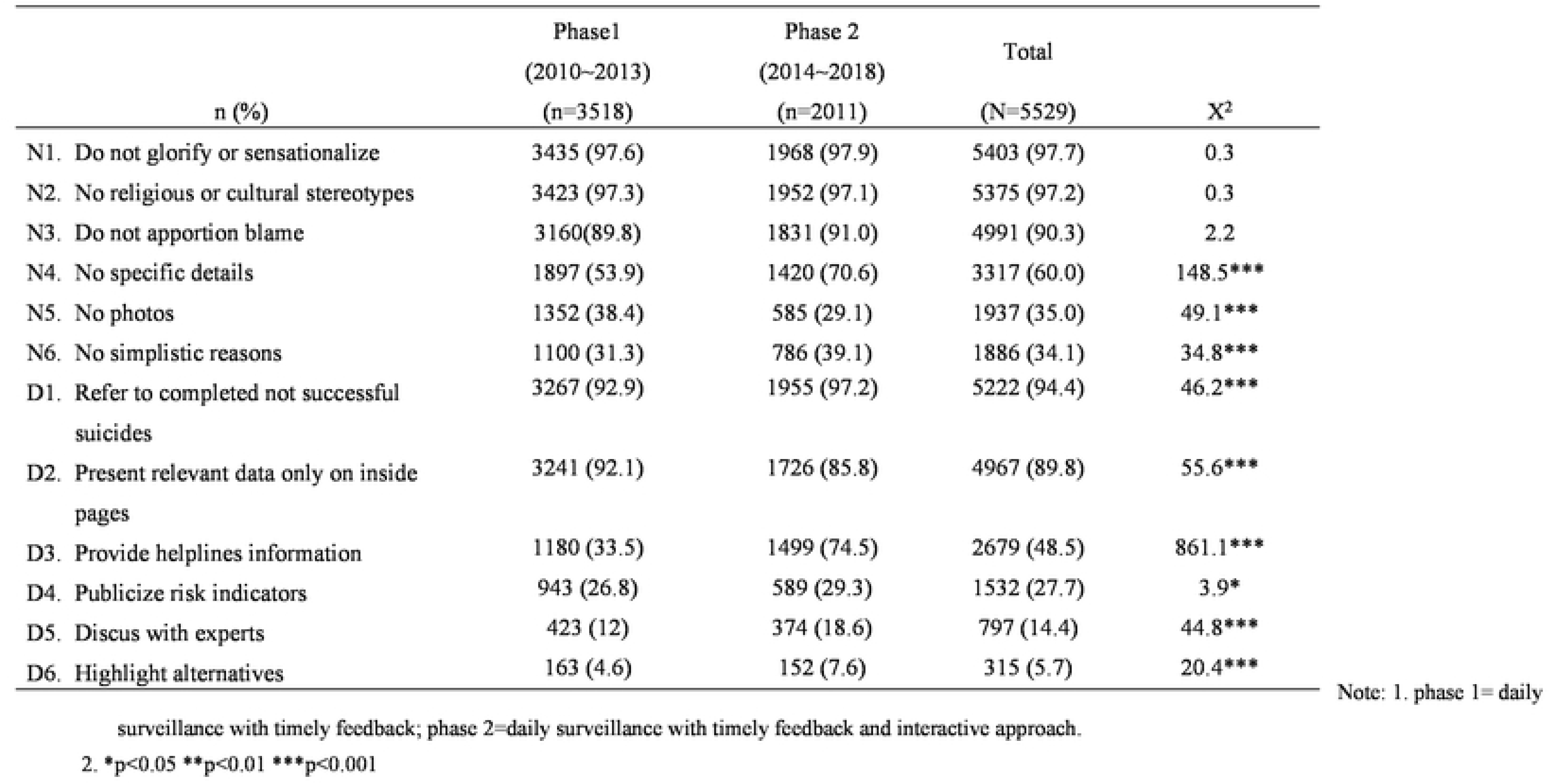
Comparison of adherence to WHO recommended guidelines between two phases.

### Adherence to WHO guidelines for online media

With respect to the online suicide reporting during 2017-2018 (Table 3), the patterns of adherence profiles for the six Dos and Don’ts by three-month intervals over the two-year observation period were similar to those of the print media. For the Don’ts, the items “Do not glorify or sensationalize,” “Do not apportion blame” and “No religious or cultural stereotypes” had the best adherence (close to 100%); the item “Do not report specific details of the suicide method” continuously improved in adherence from 82.6% to 89.9% during the two-year observation period. As in print, the items “No photos or suicide notes” (average 21.5%) and “No simplistic suicide reasoning” (average 36.4%) had lower levels of adherence (Supplementary Figure 4).

**Table 3:** The bimonthly rates of adherence to WHO recommended guides for online newspapers during 2017 and 2018

| Month/year<br>n (%) | 4~6/2017<br>(n=979) | 7~9/2017<br>(n=1043) | 10~12/2017<br>(n=989) | 1~3/2018<br>(n=755) | 4~6/2018<br>(n=1112) | 7~9/2018<br>(n=770) | 10~12/2018<br>(n=666) | Total<br>(N=6314) | X<sup>2</sup> |
| --- | --- | --- | --- | --- | --- | --- | --- | --- | --- |
| N1. Do not glorify/ sensationalize | 973 (99.4) | 1040(99.7) | 982 (99.3) | 752 (99.6) | 1030(92.6) | 737 (95.7) | 644 (96.7) | 6158 (97.5) | 183.9*** |
| N2. No religious or cultural stereotypes | 970 (99.1) | 1030(98.8) | 982 (99.3) | 749 (99.2) | 1102(99.1) | 760(98.7) | 661 (99.2) | 6254 (99) | 3.1 |
| N3. Do not apportion blame | 970 (99.1) | 1032(98.9) | 980 (99.1) | 751 (99.5) | 1093(98.3) | 764 (99.2) | 664 (99.7) | 6254 (99) | 11.4 |
| N4. No specific details | 809 (82.6) | 893 (85.6) | 808 (81.7) | 658 (87.2) | 1000(89.9) | 681 (88.4) | 588 (88.3) | 5437 (86.1) | 46.6*** |
| N5. No simplistic reasons | 370 (37.8) | 327 (31.4) | 219 (22.1) | 234 (31) | 487 (43.8) | 405 (52.6) | 259 (38.9) | 2301 (36.4) | 223.9*** |
| N6. No photos | 299 (30.5) | 242 (23.2) | 122 (12.3) | 153 (20.3) | 277 (24.9) | 149 (19.4) | 117 (17.6) | 1359 (21.5) | 114.9*** |
| D1. Refer to completed not successful suicides | 979 (100) | 1042(99.9) | 987 (99.8) | 753 (99.7) | 1011(90.9) | 760 (98.7) | 666 (100) | 6198 (98.2) | 399*** |
| D2. Present relevant data only on inside pages | 916 (93.6) | 972 (93.2) | 925 (93.5) | 731 (96.8) | 1040(93.5) | 755 (98.1) | 661 (99.2) | 6000 (95) | 67*** |
| D3. Provide helplines information | 890 (90.9) | 942 (90.3) | 899 (90.9) | 698 (92.5) | 948 (85.3) | 710 (92.2) | 629 (94.4) | 5716 (90.5) | 53.9*** |
| D4. Publicize risk indicators | 52(5.3) | 87 (8.3) | 100 (10.1) | 104 (13.8) | 104 (9.4) | 38 (408) | 50 (7.5) | 535 (8.5) | 57.6*** |
| D5. Discuss with experts | 27 (2.8) | 18 (1.7) | 14 (1.4) | 14 (1.9) | 19 (1.7) | 8 (1) | 4 (0.6) | 104 (1.6) | 14.3* |
| D6. Highlight alternatives | 190 (19.4) | 170 (16.3) | 197 (19.9) | 112 (14.8) | 232 (20.9) | 167 (21.7) | 214 (32.1) | 1282 (20.3) | 83.5*** |

\*p<0.05 \*\*p<0.01 \*\*\*p<0.001

With regard to the Dos, the three items of “Refer to completed, not successful suicides,” “Publish relevant data only on inside pages” and “Provide helpline information” had the steadily highest rates of adherence (higher than 95%) across two years. As with the print media, poor adherence was found for items of discussing with experts (2.7%), publicizing risk indicators (21.1%) and highlighting alternatives (22.2%). Compared to print media (Table 4), the online media had similar trends in adherence to the 12 recommendations, but significantly lower compliance for the items “No photographs,” “No simplistic reasoning,” “Discuss with expert” and “Publicize risk indicators” (Supplementary Figure 5).

**Table 4:** Comparison of adherence to WHO guidelines between online and print newspapers in 2017-2018

| n (%) | Online (N=6314) | Print (N=568) | p-value* |
| --- | --- | --- | --- |
| N1. Do not glorify or sensationalize | 6158 (97.5) | 564 (99.3) | 0.007 |
| N2. No religious or cultural stereotypes | 6254 (99) | 555 (97.7) | 0.003 |
| N3. Do not apportion blame | 6254 (99) | 526 (92.6) | <0.001 |
| N4. No specific details | 5437 (86.1) | 461 (82.2) | 0.001 |
| N5. No photos | 1359 (21.5) | 153 (26.9) | 0.003 |
| N6. No simplistic reasons | 2301 (36.4) | 295 (51.9) | <0.001 |
| D1. Refer to completed not successful suicides | 6198 (98.2) | 559 (98.4) | 0.66 |
| D2. Present relevant data only on inside pages | 6000 (95) | 525 (92.4) | 0.007 |
| D3. Provide helplines information | 5716 (90.5) | 480 (84.5) | <0.001 |
| D4. Publicize risk indicators | 535 (8.5) | 85 (15.0) | <0.001 |
| D5. Discuss with experts | 104 (1.6) | 91 (16.0) | <0.001 |
| D6. Highlight alternatives | 1282 (20.3) | 50 (8.8) | <0.001 |

\* The p value was derived from the Chi-squared test.

## Discussion

Newspaper reports have significant influence among various media and represent an important source of suicide coverage for readers. Most online reporting materials are based on the content of print reporting. The present study demonstrated that the suicide reports in print newspapers apparently decreased over the nine years between 2010 and 2018. However, the number of online items covering suicide increased dramatically. This difference indicates that suicide coverage by number of articles reported in print media could be more effectively and easily controlled than the online media under a policy of self-regulation and routine government monitoring. However, online news media apparently will have more suicide reporting because of its strengths of no space limit, immediate release of news and rapid dissemination.

In spite of the wide dissemination and promotion of WHO guidelines for media suicide reporting, most studies report a generally low reported adherence to guidelines and show cultural differences in compliance with various guideline items [16-18, 19-24]. For instance, some countries reported a quite low adherence to providing helpful information and higher adherence with not publishing photographs. Most importantly, few cohort studies have examined the feasibility of guideline implementation and the effectiveness of a long-term intervention in terms of adherence to media guidelines by item and by type of media (print or online). The results of the present study demonstrated that the long-term trends of adherence in both print newspaper and online media reporting over nine years could be categorized into three patterns: 1) sustained higher adherence, 2) sustained lower adherence and 3) steadily marked improvement. Sustained lower adherence was found in one Don’t (“Don’t publish photographs or suicide notes”) and three Dos (“Highlight alternatives to suicide,” “Discuss with experts” and “Publicize risk indicators and warning signs”). Adherence of these four items fluctuated and showed no obvious continuous improvement over the years. The first recommendation against photography is in general conflict with the media’s intent of using pictures to attract the reader’s attention. The latter three items are related to mental health literacy, which may be low in journalists; close collaboration between media professionals and mental health authorities could improve adherence to these recommendations. In addition, it is not easy for the media to quickly access mental health experts after a suicide event or to report risk signs or alternatives in a single report which may have space limitations.

Because photographs produce a strong impact on readers [14], violation of publishing photographs was apparently the most serious issue in Taiwan. In this study, we used strict criteria to rate all photographs as noncompliant with the guidelines. Much lower adherence to the “No photograph” recommendation (about 9%) was also reported by Fu et al [16]. Chiang et al. found that suicide was commonly reported on the front page with photographs [17].

During our surveillance period, the media could withdraw immediately the inappropriate online photos upon our advice, but publishing photographs remains an issue. Taiwanese media fully accept that publishing actual pictures of the deceased body or persons is absolutely inappropriate and should be avoided. However, there is room for discussion about the relative contraindication for presentation of an associated photograph, graphic or image. For instance, they could absolutely respect the suicide people’s privacy and dignity by withholding photographs, but then use a modified illustrating diagram or animation that implicitly includes suicide details (e.g., location or method), without sensitivity to the possible negative impact in terms of the copycat effect (also called the Werther effect). Despite years of evidence-based studies and enhanced media-health professional communication, we have been unable to convince news outlets of the possible imitative effect on suicide behavior and stop them publishing photographs. With recommendations from the TSPC, the Taiwan legislative Yuan recently passed a Suicide Prevention Act on May 31, 2019 that clearly prohibits the photographic or image presentation of suicide persons or places, as well as a detailed description of the suicide scene or method. Continuous communication and legal restriction are expected to be effective in preventing the future presentation of inappropriate photographs and images.

### Effectiveness of TSPC surveillance

Regarding the effectiveness of TSPC surveillance with the interactive intervention, there was a significant increase in adherence rate for all items except for “No photograph,” based on the trend analysis (Table 1) and two-phase comparison (Table 2). In particular, continuous and markedly increased adherence by year was noted for three items: “Do not report the details of suicide methods,” “No simplistic reasons” and “Provide information on helplines and community resources.” A stable trend of very good adherence was maintained over nine years for the items “Refer to suicide as completed, not successful” and “Publish relevant data only on inside pages.” These items are related to concepts that are well accepted and feasible on the media side. Most importantly, unlike reports from most other countries, both print and online newspapers demonstrated the greatest improvement in providing information on toll free helplines. This long-lasting good adherence shows that our intervention had positive effects and that the media has matured in its reporting style. However, even with statistically significant improvement, many articles still had lower adherence to “Work closely with experts or health authorities,” and “Provide alternatives to suicide” and “Publicize risk indicators.” In reality, the majority of suicide news reporting occurs in an emergent situation, with health experts not always available to respond to the events. To resolve this problem, the TSPC quickly submits special articles to the media on related risk factors or coping alternatives, as a complementary balance to insufficient or inadequate suicide reporting. Our long-term observation strongly suggested that some of the “Do” items are important for responsible reporting, but are not the journalist’s duty only; they could be fulfilled by adjunctive articles, written either by health professionals or journalists in other sections as an integrated part of the suicide coverage. Therefore, effective suicide prevention will need to increase the practicality of the guidelines for media professionals and promote the collaboration of multidisciplinary professionals involved in this area.

The role of the media in suicide prevention and health promotion is well recognized. The adherence of online media to the WHO suicide reporting guideline has become a bigger issue because of its quick dissemination and transmission by internet users. Further guidelines are needed, not only for media personnel, but also for the general public. It is necessary to constantly emphasize the media’s social responsibility as an important health gatekeeper. Our study of long-term daily monitoring and evaluation of the observance of the WHO guidelines for suicide reporting demonstrated the effectiveness of such monitoring and the limitations to further quality improvement in suicide reporting. This area promises to develop into a new specialty of liaison between the media and psychiatry. Developments at the interface of media reporting and mental health can yield public health benefits in terms of suicide prevention and improved mental health literacy.

### Limitations

The study focused only on major newspapers and did not involve other domestic newspapers or other forms of media such as social networking media on the internet. The results therefore represent only the mainstream news reporting styles on suicide. We also did not survey the general public for the possible negative impact of poor quality reporting or analyze other potential outcomes.

## Acknowledgments

This work was supported by the Ministry of Health and Welfare, Executive Yuen, Taiwan (L.M.B., grant number M04B3020&M07B4116). The authors thank all our colleagues at the Taiwan Suicide Prevention Center for their active participation in data collection and administrative help.

